# Hidden in plain sight: Bacterial genomes reveal thousands of lytic phages with therapeutic potential

**DOI:** 10.1101/2025.05.06.652010

**Authors:** Alexander Perfilyev, Anastasiya Gæde, Steve Hooton, Sara A Zahran, Caroline Sophie Winther-Have, Panos G Kalatzis, Rodrigo Ibarra Chavez, Rachael C Wilkinson, Anisha M Thanki, Bent Petersen, Zhengjie Liu, Qing Zhang, Qianghua Lv, Yuqing Liu, Adriano Gigante, Robert J Atterbury, Andrew Millard, Martha Clokie, Thomas Sicheritz-Pontén

## Abstract

Phages are typically classified as temperate, integrating into host genomes, or lytic, replicating and killing bacteria. Lytic phages are not expected in bacterial genome sequences, yet our analysis of 3.6 million bacterial assemblies from 1,226 species revealed >100,000 complete lytic phage genomes, which we term Bacterial Assembly-associated Phage Sequences (BAPS). This represents a ∼five-fold increase in the number of phages with known hosts and fundamentally reshapes our understanding of phage biology.

Identifying BAPS has enabled the discovery of entirely novel phage clusters, including clusters distantly related to *Salmonella* Goslarviruses in *E. coli*, and *Shigella*, while significantly expanding known genera such as *Seoulvirus* (from 16 to >300 members). Notably, close relatives of therapeutic lytic phages were also detected, suggesting clinical isolate sequencing unknowingly archives viable phage candidates. The discovery of complete, lytic phage genomes within bacterial assemblies challenges long-standing assumptions and reveals a vast, untapped reservoir of phages.

## Introduction

Bacteriophages are traditionally divided by lifestyle into two categories: temperate, which integrate into bacterial chromosomes as prophages, and lytic, which replicate rapidly and kill their hosts^1^. According to this binary framework, only temperate phages are expected to persist in bacterial genome sequencing datasets, because lytic phages, in contrast, would be expected to eliminate their hosts during infection. Therefore, by definition the bacteria that are sequenced should be those that have escaped lytic predation^2^.

To contextualise the data we present, it is important to understand how bacterial isolates are typically sequenced. Generally single colonies are obtained, to ensure they are axenic and represent a clonal population and these are then grown in liquid culture then sequenced^3^.

During the process of investigating jumbo phages that infect *Salmonella* spp, we encountered something unexpected within bacterial genomes: complete, intact genomes of lytic phages embedded within publicly available bacterial genome sequences. Initially we assumed this observation to be an anomaly or assembly artefacts, but on further analysis these sequences appear repeatedly, across species, geographies, and sequencing projects.

Prompted by this observation, we carried out further analysis and systematically examined ∼3.6 million bacterial genome sequences that span 1,226 species from the NCBI RefSeq database. The results were striking. as over 100,000 complete lytic phage genomes were identified, many of which belong to previously unrecognised genera uncovered entirely new clusters of jumbo phages within *Salmonella, E. coli*, and *Shigella*. In other cases, for example in the *Seoulvirus* genus, we expanded the number of known phages from fewer than 20 to over 300 representatives.

These findings challenge the expectation that only temperate phages are retained in bacterial genome data. The presence of intact, lytic phage genomes across diverse taxa suggests that these phages were caught during a snapshot of lytic infection during the point that the bacteria are sequenced, or that they may persist within bacterial cells under conditions that suppress lysis.

One possible explanation is that the phages are present in what is termed the carrier state or pseudolysogeny which is an underexplored infection mode in which lytic phages coexist with their hosts without integrating into the chromosome or initiating replication^4^. While the mechanism remains unclear, their prevalence across taxa and geographies demands a re-evaluation of how phages are maintained and missed in sequencing data.

Here, we present a large-scale analysis of these hidden lytic phage sequences, which we term Bacterial Assembly-associated Phage Sequences (BAPS). We describe their host distribution, taxonomic diversity, and overlap with known therapeutic phages, with significant implications for both microbial ecology and the development of phage-based therapeutics.

## Results

To identify complete lytic bacteriophage genome sequences within bacterial genome assemblies (BAPS), we developed a comprehensive bioinformatic workflow, starting with assembly data available from NCBI. We filtered and analysed 3.6 million bacterial assemblies, focusing on contigs larger than 5,000 bp as potential phage candidates. Using the phage-likeliness predictor *phager*.*py* tool (Supplementary material), we refined this set to 3.5 million contigs of putative phage origin. These contigs were further screened against reference databases to identify sequences of bacteria, plasmid, and phage origin. We identified 119,510 lytic phages and clustered them using Average Nucleotide Identity (ANI) to distinguish lytic from temperate phages. This resulted in the identification of 119,510 lytic phages, 602,285 plasmids, and 146,575 temperate phages.

The distribution and abundance of BAPS contigs within all bacterial genome sequences is shown in **Figure 1**. Mapping the bacterial host for each BAPS to a reference bacterial phylogenetic tree, reveals the widespread presence of lytic phage genomes within bacterial genome assemblies of diverse origins.

**Figure 1.**
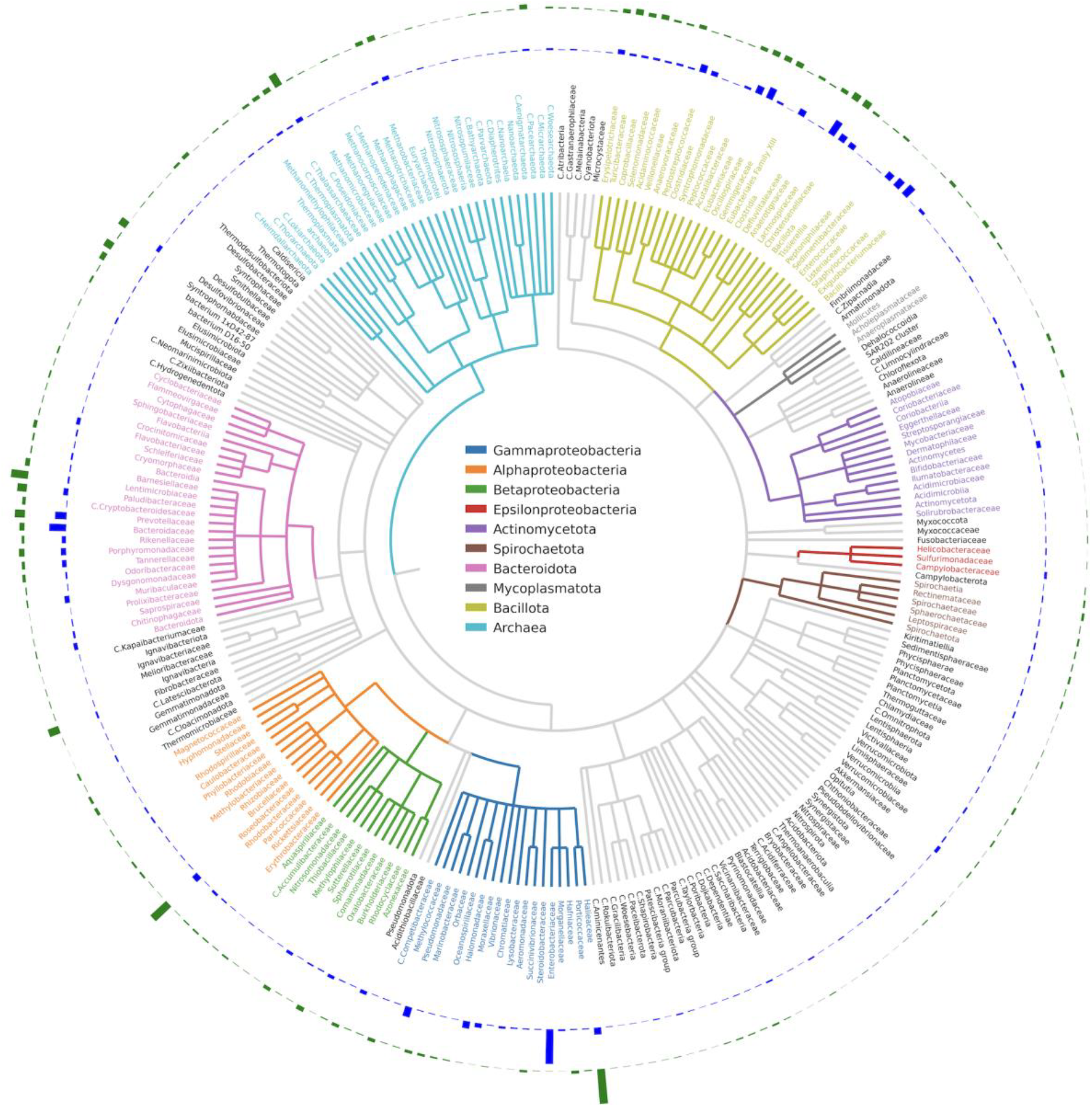
Phylogenetic distribution of BAPS across bacterial families. The circular phylogenetic tree displays bacterial families, with each bacterial class color-coded according to the legend inside the circle. Only families with at least five observed BAPS are shown. The outer blue bars indicate the number of BAPS found per bacterial family. The outermost green bars represent the ratio of BAPS to the total number of genomes available for each family.

Clearly, if BAPS are distributed evenly across bacterial taxa, the largest numbers will naturally be found within bacterial species that have been extensively sequenced, such as those targeted in clinical surveillance or outbreak investigations. To determine how sequencing bias relates to BAPS discovery, we examined the ratio of BAPS relative to the total number of genome assemblies available for each bacterial family.

The *Gammaproteobacteria* harbour the highest total number of BAPS, primarily due to the large number of BAPS contigs found within *Enterobacteriaceae* genome assemblies. This pattern reflects the intensive sequencing of pathogenic species within this family, particularly *Escherichia coli* and *Salmonella spp*., which dominate public genome datasets.

Notably, BAPS are also found at relatively high frequency within the phylum *Bacillota*. Clinically important Gram-positive families, including *Staphylococcaceae, Enterococcaceae, Streptococcaceae*, and *Clostridiaceae*, all contain BAPS contigs within their genome assemblies.

Beyond clinically relevant taxa, several environmental bacterial families also exhibit a high BAPS-to-sample ratio despite relatively limited sequencing effort. For example, within the *Alphaproteobacteria*, both the *Roseobacteraceae*, key players in marine biogeochemical cycling, and the *Acetobacteraceae*, a group of acetic acid bacteria, show a notable enrichment of BAPS.

Overall, our analysis of bacterial classes and families (**Figure 2**) highlights consistent patterns of BAPS distribution, with lytic phage genomes embedded within bacterial assemblies across diverse environments and hosts.

**Figure 2.**
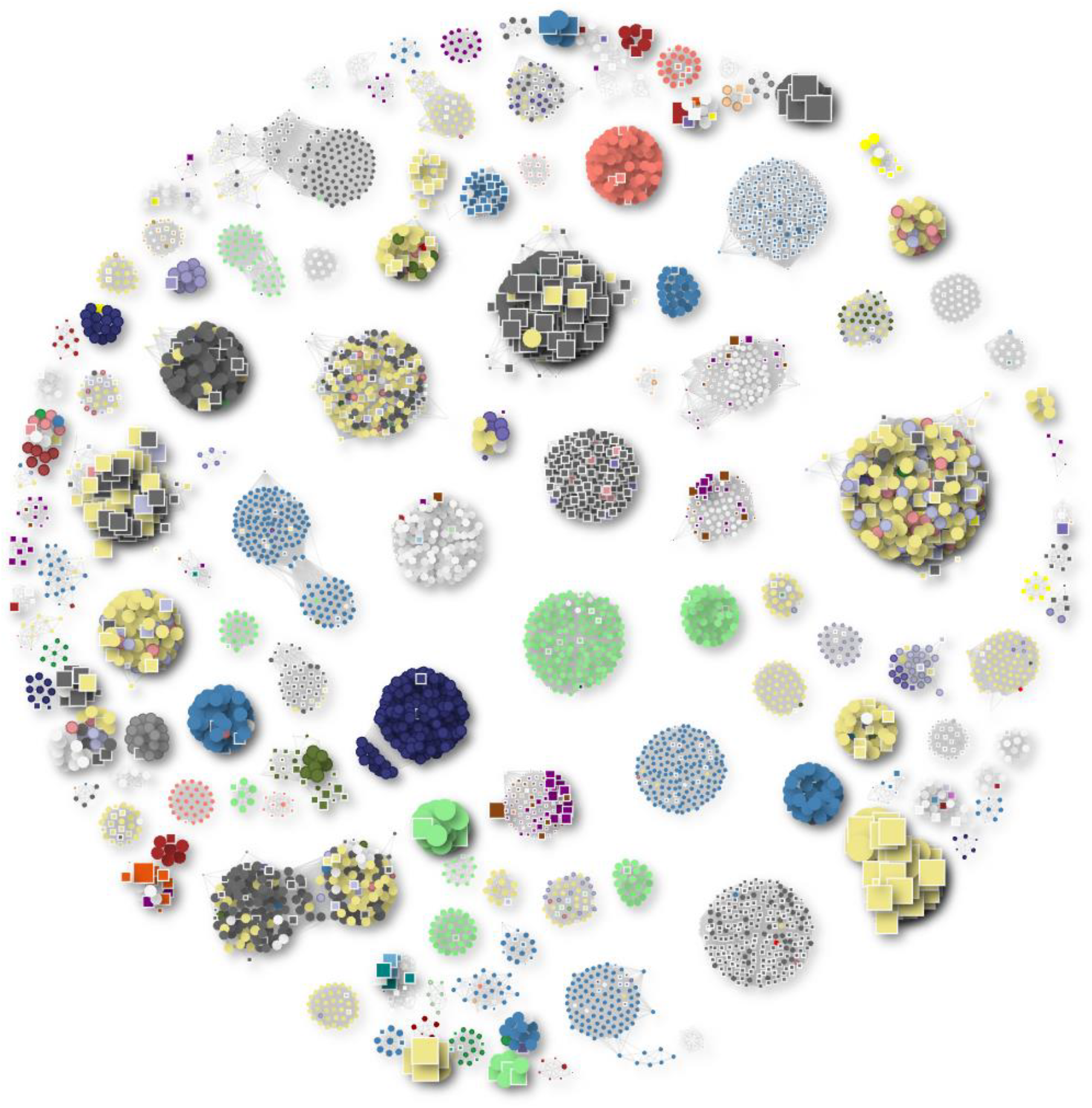
Clusters of phage genomes containing both NCBI and BAPS phages with a minimum genome size of 40 kb. Each cluster shown consists of at least five members, including both phages from the NCBI database and BAPS phages. Phage genomes are represented as circles, while BAPS genomes are depicted as squares. The size of each shape is proportional to the genome size. Clusters are color-coded based on the host genus: *Pseudomonas* (light green), *Klebsiella* (blue), *Escherichia coli* (khaki), *Salmonella* (dark grey), and *Serratia* (red).

Given the dominance of BAPS within the *Enterobacteriaceae*, we focused on BAPS associated with *Salmonella spp*. and *Escherichia coli* where we identified six distinct lineages of lytic jumbo phages. In several cases, the number of known members within a given lineage has expanded dramatically. For example, the genus *Seoulvirus* has increased from 20 reference phage genomes in GenBank to over 300 complete genomes.

Similarly, the orphan jumbo phage genus *Goslarvirus*, originally represented only by the phage Goslar^5,6^, has been expanded from 1 to 237 genomes in our dataset.

Importantly, our approach has also led to the discovery of a novel jumbo phage genus, for which we propose the name “Bapsvirus”. In total, we identified 247 BAPS genomes within this cluster, with the largest 54 genomes each approximately 220 kb in size. These phages are associated with *Salmonella* spp., *E. coli*, and *Shigella* spp., illustrating that bacterial genome assemblies represent a valuable and untapped resource for phage discovery.

### Expansion of existing Salmonella jumbo phage groups

#### *Seoulvirus* — Massive Expansion of a Therapeutically Relevant Phage Genus

The largest BAPS cluster identified within *Salmonella* and *E. coli* genome assemblies belongs to the genus *Seoulvirus* in the family *Chimalliviridae*. We identified over 300 new *Seoulvirus* jumbo phage genomes (239–242 kb), expanding the known diversity of this group by more than an order of magnitude (**Figure 3A**). These phages are well-characterised lytic viruses with documented therapeutic potential against *Salmonella spp*.^7^ The widespread detection of *Seoulvirus* BAPS across human, animal, and environmental isolates underscores a stable and pervasive host-phage association in diverse environments.

**Figure 3.**
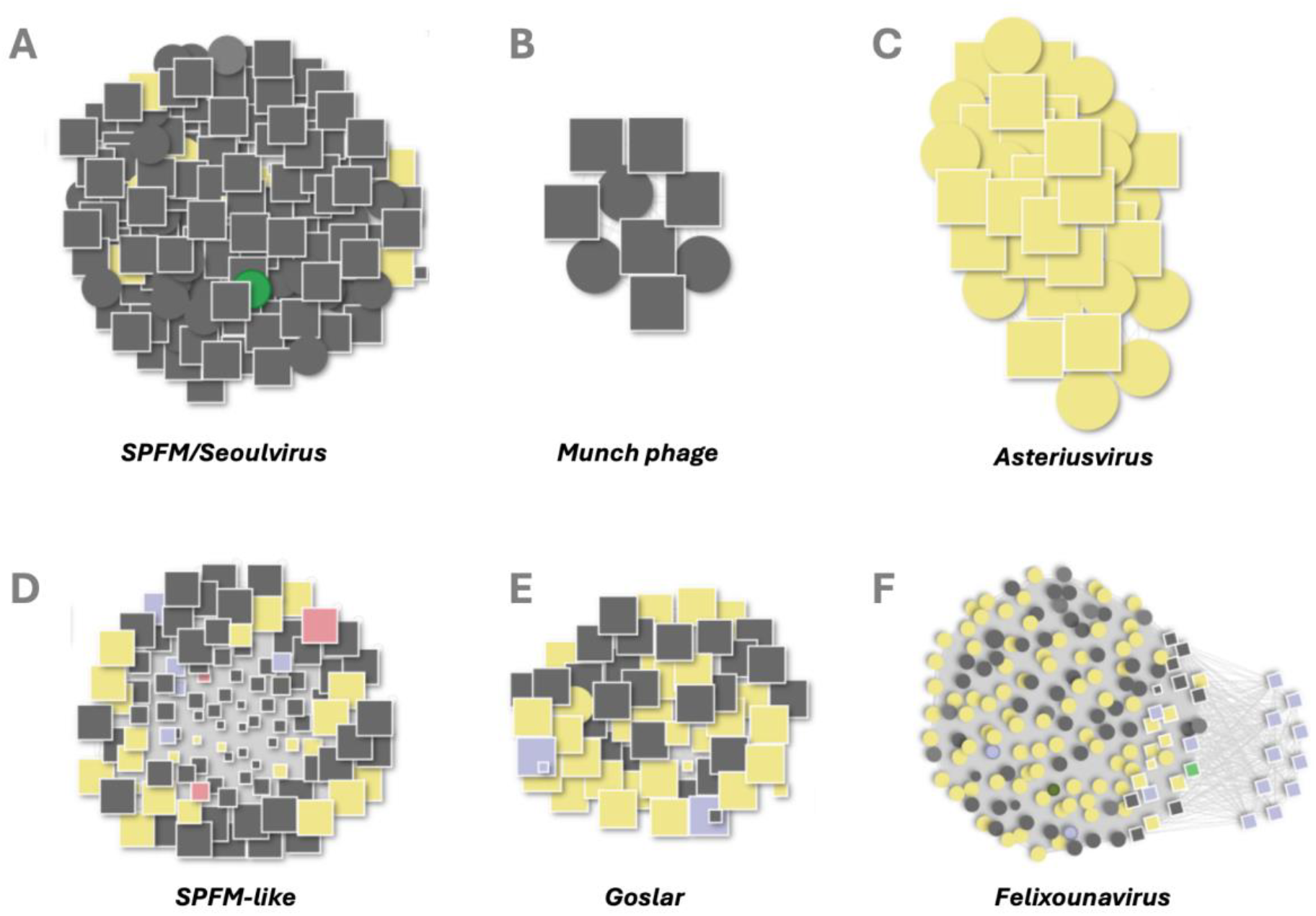
Clusters of phages associated with *Salmonella* and related genera. Each circle represents a known phage genome from NCBI, while each square represents a BAPS-identified genome. The size of each shape is proportional to genome length. The color indicates the bacterial host genus: grey = *Salmonella spp*, yellow = *Escherichia coli*, green = *Clostridium spp* (likely a misannotation of the bacterial sequence), and pink = *Shigella spp*. Phage and BAPS genomes are connected if they share a Mash distance ≤ 0.1. The clusters highlight distinct phage lineages, including Goslar, Bapsvirus, Munch phage, HeyDay, Asteriusvirus, and Seoulvirus.

#### “Bapsvirus” — Discovery of a Novel Jumbo Phage Genus in *Salmonella*

Excitingly, our approach has also identified a distinct cluster of 247 BAPS genomes (∼220– 249 kb), representing a novel jumbo phage genus, for which we propose the name “Bapsvirus” (**Figure 3D**). These phages show limited sequence similarity (15–40% ANI) to *Seoulvirus*, but retain conserved genome architecture and synteny. Most BAPS contigs were found in *Salmonella* genome assemblies, with additional sequences in *E. coli* and *Shigella*. Phylogenetic analysis supports the classification of “Bapsvirus” as a new genus within the family *Chimalliviridae*, targeting bacterial cells of the *Enterobacteriaceae*. Further analysis of high-quality genomes within these two genera using *taxMyPhage*^*8*^, expanded the number of species in *Seoulvirus* from one to six, and the creation of 32 new species in the genus “Bapsvirus”.

#### “Munchvirus” — A Widely Distributed Phage Genus in *Salmonella*

BAPS analysis uncovered several additional jumbo phages related to the rare 350 kb phage Munch, tripling its known relatives and spanning diverse Salmonella isolates (**Figure 3B**). Together with known phages Munch, PHA46, SE-PL, 7t3 they form a single genus based on *taxMyPhage* analysis. We propose they are classified into a new genus “Munchvirus”,after the first isolate. BAPS analysis highlights phages of the genus “Munchvirus” are globally distributed but have remained under-sampled prior to this work.

#### *Asteriusvirus* — Expansion of an *E. coli* Jumbo Phage Genus

We identified 189 BAPS related to *Asteriusvirus*, expanding this group of *E. coli*-infecting jumbo phages (350–380 kb, ∼34% GC) from a few genomes to a much larger dataset (**Figure 3C**). Taxonomic classification increased the number of species from two to 14, and revealed a new related genus containing four species, and confirmed by phylogenetic analysis. We propose the genus name “Lethbridgevirus” after the submitting organisation. Those were identified across *E. coli* genome assemblies from diverse geographical regions and environments, confirming that *Asteriusvirus* phages are globally distributed, with the newly identified Lethbridgevirus capable of infecting both *E. coli* and *Salmonella*.

#### Felixounavirus

We identified 114 BAPS contigs related to phages in the genus *Felixounavirus*, a well studied group of phages that have been suggested to be useful for biocontrol of *Salmonella*^*9*^

#### Goslar phage — a globally distributed lytic jumbo phage with variable abundance in bacterial genome assemblies

To test whether phage contigs could be identified using a reference genome outside the *Salmonella* phage sequence space, we selected the orphan *E. coli* phage vB_EcoM_Goslar. BlastN searches confirmed that Goslar has no close relatives among known phages, however, using the BAPS pipeline, we identified 237 contigs matching *Goslarvirus*. These contigs represent a globally distributed group of lytic phages infecting pathogenic Gram-negative *Enterobacteriaceae* (**Figure 3E**), recovered from diverse environments and hosts, including water, humans, cattle, pigs, chickens, and bonobos, across diverse geographic regions. *Goslarvirus* sequences were associated with a wide range of *E. coli* serotypes and pathotypes and expanded into multiple *Salmonella* serovars and *Shigella* species. Taxonomic classification expanded the number of species, from one to 38.

The widespread recovery of *Goslarvirus* contigs across such diverse hosts and environments highlights the broad ecological success of this previously unrecognised lytic phage lineage. To further characterise these phages and assess their potential impact on bacterial genome assemblies, we examined the relative abundance of *Goslarvirus* genomes within their respective sequencing datasets.

Read mapping of 55 representative assemblies revealed striking variation in the proportion of reads mapping to phage versus host genomes (**Figure 4**).

**Figure 4.**
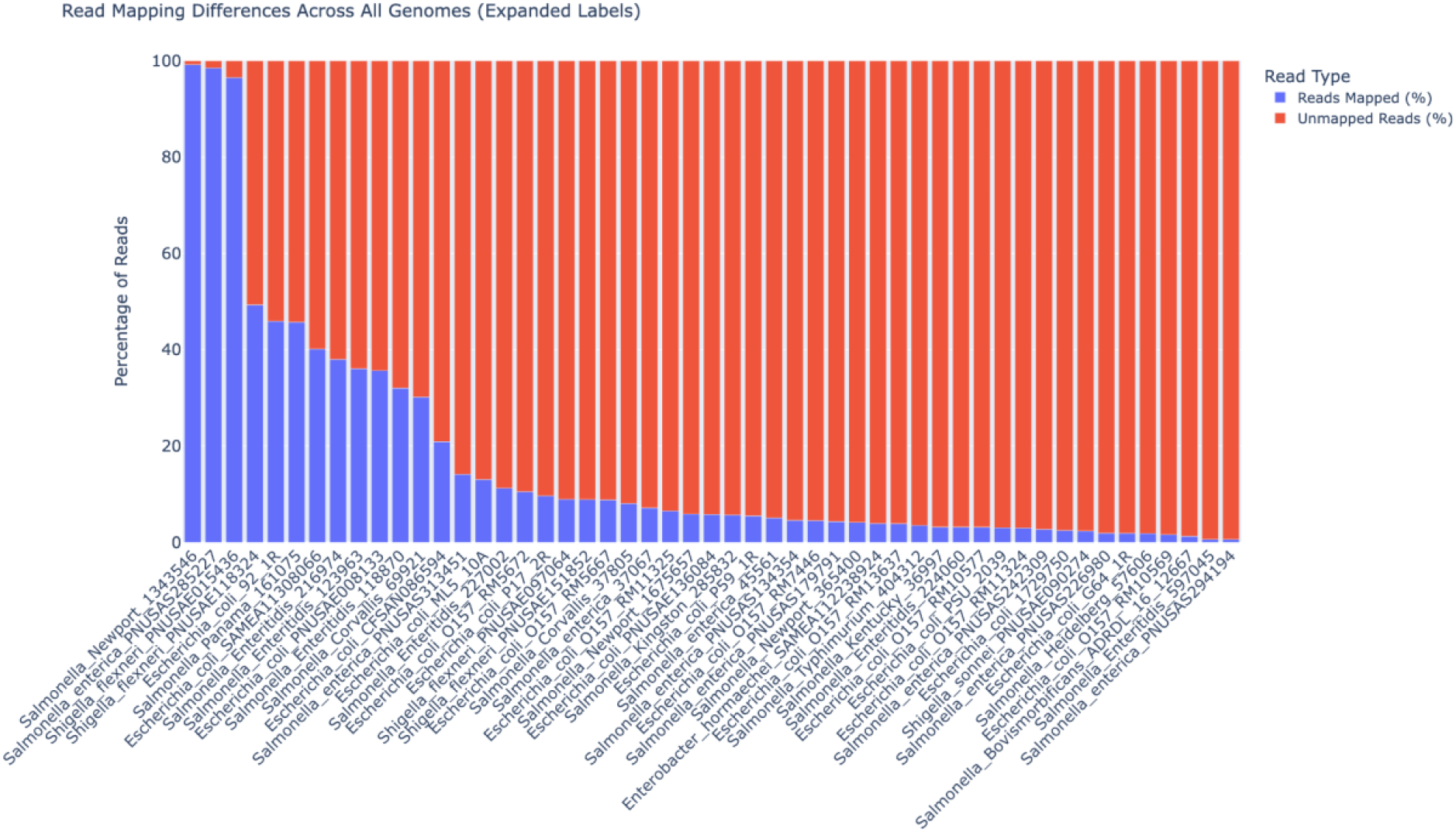
Variable abundance of *Goslarvirus* sequences in bacterial genome assemblies. Percentage of reads mapping to *Goslarvirus* contigs (blue) or remaining unmapped (red) across 55 bacterial genome assemblies. Read sets were retrieved from the NCBI SRA and mapped using Bowtie2. Phage read abundance varied markedly between samples, reflecting differences in phage load at the time of sequencing.

In some cases, *Goslarvirus* reads dominated the dataset, consistent with high-titre phage presence at the time of DNA extraction, whereas in others, phage sequences were present at very low levels. For example, in *Salmonella* Newport 134356, 99.2% of reads mapped to a 237 kb *Goslarvirus* genome, while only 0.8% mapped to the bacterial chromosome. In contrast, other assemblies showed very low phage representation, such as *Salmonella enterica* PNUSA294194, where only 0.6% of reads mapped to a 239 kb *Goslarvirus* genome. Intermediate cases were also observed, such as *Shigella flexneri* PNUSAE118324, where reads were nearly evenly split between host (50.7%) and phage (49.3%).

This variation likely reflects differences in infection dynamics, contamination, or DNA extraction protocols that favour phage particles. These findings underscore the need to consider phage content when interpreting bacterial genome data, particularly in clinical or surveillance settings where high phage abundance may influence assembly quality and downstream analyses.

### Phage Therapy Connections and Human Microbiome Relevance

It is interesting to speculate if BAPS can provide useful information on the behaviour and relationships of phages that are used during therapy and also if BAPS could represent novel sources of therapeutically relevant phages. To answer this we looked to see if known therapeutically relevant phages had BAPS homologues. The presence of therapeutically-related phages in BAPS would support the idea that human and animal exposure to these phages is part of natural bacterial dynamics. It may also have relevance to the possible presence of neutralising antibodies for these particular phages within the human/animal body.

To evaluate this relationship of BAPS, we examined their similarity to 66 therapeutic phages with publicly available genomes (Sup Table 3). These are all lytic phages that were previously used in human or animal therapy. Of note, 55 showed at least one BAPS match with moderate genetic similarity (roughly corresponding to ≥ 80% ANI), and 39 had highly similar counterparts (≥ 95% ANI).

Several BAPS equivalents were seen in 18 distinct clusters of therapeutically relevant phages (**Figure 5**). Notably, a large *Escherichia coli phage* cluster containing phage T4^10^, included phages used in clinical or animal studies by Bruttin & Brüssow (2005), Guo et al. (2021), and Pirnay et al. (2024).

**Figure 5.**
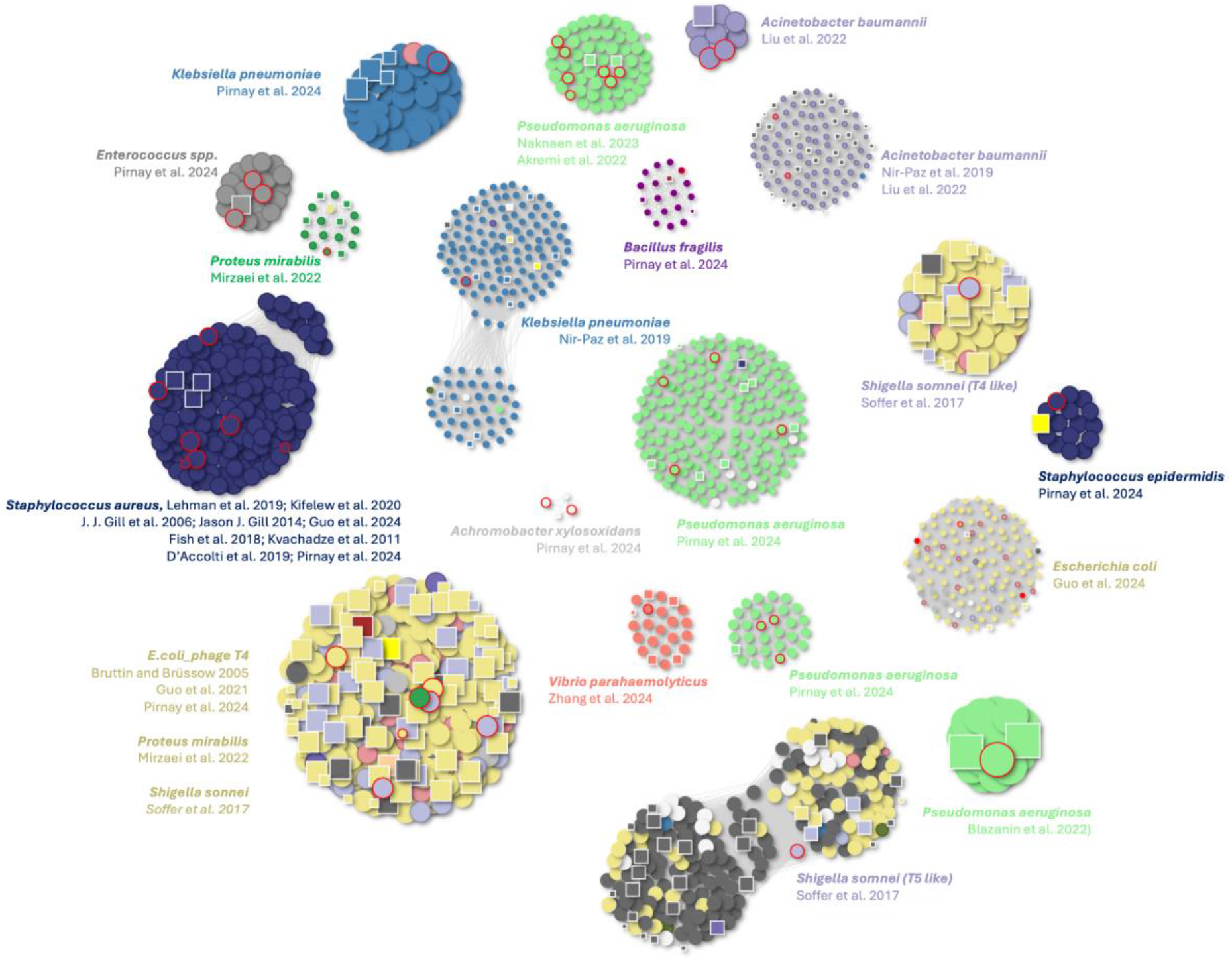
Clusters of lytic phages used in phage therapy and their genomic relationships to BAPS-derived phages. Each node represents a phage genome: circles denote lytic phages from Genbank, and squares represent lytic phages discovered within BAPS bacterial assemblies. Nodes outlined in red indicate phages that have been used in published phage therapy studies. Edges connect genomes with a Mash distance ≤ 0.1, indicating high sequence similarity. Colours represent host genera.

Similar patterns of BAPS similarity were observed for other therapeutically relevant phages. This included *Shigella sonnei* Mosigivirusese and Tequatroviruses^11^, *Proteus mirabilis* phages^12^, and multiple *Pseudomonas aeruginosa* clusters, such as OMKO1 and PA1Ø. A large cluster associated with *Staphylococcus aureus* comprised phages used in therapy studies^13–15^ and others, many of which had highly similar BAPS counterparts, indicating that these therapeutic phages are naturally represented within the BAPS dataset. Additionally, two *Klebsiella pneumoniae* phages previously employed in human therapeutic interventions^16,17^ also demonstrated high sequence similarity to BAPS.

Multiple BAPS contigs are found associated with significant human pathogenic bacteria, including *Acinetobacter baumannii, Pseudomonas aeruginosa, Proteus mirabilis*, and *Klebsiella pneumoniae*. Phages used to target the opportunistic pathogen *Bacteroides fragilis* and the cystic fibrosis-associated *Achromobacter xylosoxidans* were also found as BAPS in a limited number of bacterial assemblies. BAPS with high similarity to phages previously employed against Gram-positive opportunists such as *Staphylococcus aureus* and *Staphylococcus epidermidis* were also observed. Some phage genomes, such as *Listeria monocytogenes* phage P100 (a component of the commercial product Listex P1007), did not match any BAPS contigs.

Although the most highly populated BAPS cluster corresponded to *P. aeruginosa* phage PA10 (n = 3146), a lytic variant of the temperate phage D3112, no further analysis was conducted on this phage due to its temperate origin, which makes its presence in bacterial assemblies expected.

We also investigated whether BAPS phages were detectable in the human microbiome **(Figure 6)**. Across four major gut virome datasets (MGV, GPD, ELGV, and GVDv1) and one whole metagenome dataset (HMP), nearly 2000 BAPS contigs were identified, with high ANI >96% across most hits. Although the number of matched metagenomic contigs varied between databases, several BAPS phages appeared consistently across multiple catalogues, supporting their ecological relevance in human microbiomes. Notably, many of these matched contigs were in the 230–240 kb size range, consistent with jumbo lytic phages.

**Figure 6.**
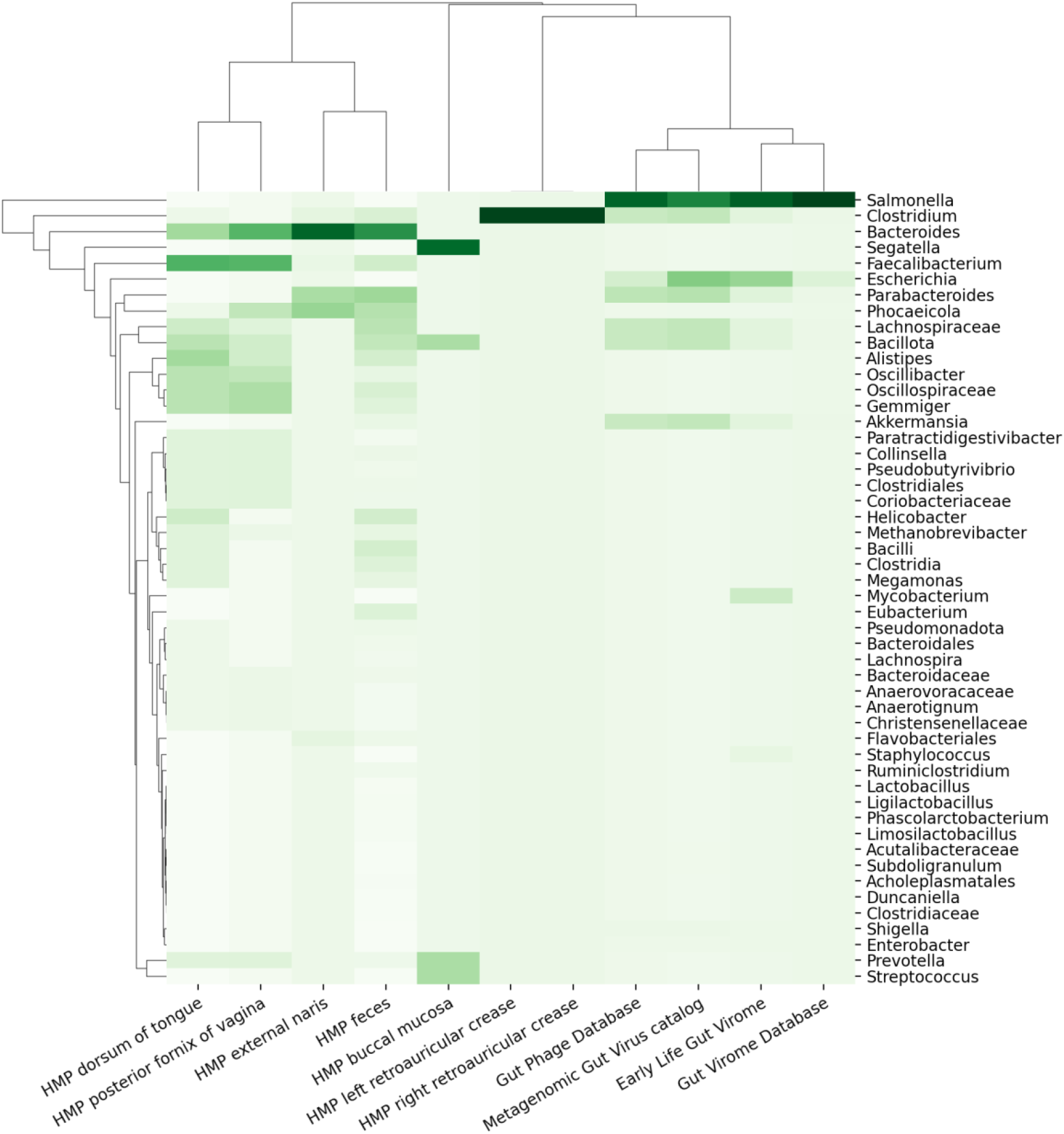
Detection of BAPS phages in human microbiome datasets. Clustered heatmap showing the distribution of BAPS phage contigs detected across various human-associated metagenomic datasets, including the Human Microbiome Project (HMP), Metagenomic Gut Virome (MGV), Gut Phage Database (GPD), Early Life Gut Virome (ELGV), and Gut Virome Database v1 (GVDv1). Columns represent different datasets, while rows indicate the top 50 bacterial genera associated with BAPS phages based on taxonomic annotation of matched contigs.

Together, these findings suggest that BAPS are present in clinical and environmental bacterial isolates and may also persist in human-associated microbial communities which motivates future work to try and establish what they may be doing.

While most BAPS-derived assemblies matched the expected bacterial taxonomy, we identified a small subset of assemblies where the taxonomic assignment did not correspond to the annotated host species. These misclassifications can lead to seemingly implausible phage-host associations. For instance, BAPS identified 11 complete lytic phage genomes within *Legionella* assemblies—an exciting prospect given the current lack of known lytic phages for this pathogen. However, our taxonomic validation pipeline (see Methods) revealed inconsistencies in these cases, suggesting that the actual bacterial source of the assemblies may not have been *Legionella*. A similar example appears in **Figure 5**, where one BAPS genome originally annotated as *Clostridium perfringens* clusters tightly with known *Staphylococcus* phages. Detailed inspection confirmed that the underlying contigs were in fact of *Staphylococcus* origin.

## Discussion

### Phage Lifecycle Complexity and the Carrier State

Phages have traditionally been classified as either virulent, directly lysing their bacterial hosts; temperate, integrating into host genomes and replicating as prophages; or chronic, continuously producing and releasing new virions without killing the host. However, the boundaries between these lifecycles are increasingly recognised as fluid, representing a continuum of phage–host interactions rather than a strict dichotomy. Several studies have highlighted this grey zone, identifying infection states such as the carrier state and pseudolysogeny, where phage DNA persists in host cells without triggering lytic replication or integration into the chromosome^4,18,19^.

In these non-committal states, phages can exist as extrachromosomal elements without initiating structural protein production or genome replication. Such strategies may represent evolutionary adaptations to suboptimal conditions, for example, during the infection of metabolically inactive cells, under nutrient limitation or low bacterial host numbers, where neither lytic nor strict lysogeny is optimal. Factors known to drive the carrier state condition are the prophage content of the bacterial cell, nutrient status, pH, and temperature^4^.

The carrier state may serve as a survival strategy for lytic phages in natural environments although to date it has only been described for temperate phages. The observations described here suggest lytic phages may have this quasi-stable relationship between phage and host.

### Implications of Lytic Phages in Bacterial Assemblies

The identification of complete lytic phage genomes within bacterial genome assemblies (BAPS) fundamentally challenges the assumption that only temperate phages are retained in such datasets. Traditionally, phages found alongside bacterial chromosomes were presumed to be prophages - latent and potentially inducible. However, the lytic phage genomes we identified were not integrated into the host chromosome but assembled alongside bacterial contigs in the same genome projects. Our analysis of 3.6 million RefSeq assemblies, spanning over 1,200 bacterial species, revealed more than 100,000 complete lytic phage genomes, which both expand known families and represent entirely novel genera.

These findings support the concept that lytic phages can persist within bacterial cells without completing a productive replication cycle. One plausible mechanism for this is that lytic phages may enter a carrier-like state within bacteria, aided by host factors that inhibit replication or assembly. It may also be that bacterial hosts withhold nucleotide precursors or trigger abortive infection pathways, stalling phage replication and particle assembly without leading to host death. Overall, our findings call for a re-evaluation of phage lifecycle definitions and suggest that lytic phages may engage in more nuanced interactions with their hosts than previously appreciated.

### Expansion of the of the family *Chimalliviridae*

The BAPS dataset provides insight into the biology and diversity of phages within the family *Chimalliviridae*. Members of the family are all Jumbo phages and produce a nucleus-like shell built from a chimallin protein which physically shields the phage DNA from host defense systems. The nucleus-like shell is rotated and centred within the cell cytoplasm by tubulin-like PhuZ filaments that assemble dynamically^6^. PhuZ is not a core component of phages of the *Chimalliviridae*, several of which have lost the gene encoding this protein. The loss of PhuZ can be sporadic, with close relatives maintaining *phuZ* or completely loss from a genus e.g. *Erskinevirus*^*20*^. Here we vastly expand the number of phages within the *Chimalliviridae*, particularly in the genus *Seoulvirus* and identify the new genus “Bapsvirus”, which also lacks homologues of *phuZ* .

The evolution of *Erskineviruses, Seoulviruses* and Bapsviruses from a single common ancestor suggests they have similar infection strategies, potentially involving sequestration or absence of nucleic acid-degrading enzymes. Phage Asesino (genus *Erskinevirus)* does not degrade host DNA during infection ^21^. If phages in the *Seoulvirueses* and “Bapsviruses” employ a similar strategy and maintain a “carrier-state” it would explain their high prevalence in bacterial genome assemblies. The specific evolutionary adaptations that enable frequent carrier state formation remain unclear. These associations deserve further investigation, as they may represent phages with particularly good therapeutic potential due to their capacity for stable, non-destructive intracellular persistence.

The BAPS dataset vastly expands the number of jumbo phage genomes associated with *Escherichia* spp, *Salmonella* spp and *Shigella* spp. Prior to this study only 49 jumbo phages infecting *Escherichia* spp and *Salmonella* spp had been reported (Feb 2025) ^22^. The BAPS pipeline expanded this to >800 genomes and identified the first to infect *Shigella* spp. This raises the question as to why they are so rarely isolated, if they can be identified as infecting bacteria. One possible explanation is these lytic phages are entering a carrier state, under the conditions of isolation that prevents clear plaque formation during the isolation process. Further testing of culture conditions may well increase the recovery of jumbophages during isolation-based approaches.

### Implications for Phage Therapy

The unexpected discovery of lytic phages within bacterial assemblies offers new possibilities to interpret and develop therapeutic phages. The presence of known therapeutic phages in bacterial genomes indicates that they are naturally occurring and potentially adaptable for therapeutic use. This could enhance the safety profiles of phages used in therapy, as these phages have already demonstrated the ability to coexist with their bacterial hosts. The regulatory framework for phage therapy could be revised in light of these findings.

Important questions also arise from our observations especially when considering phage therapeutic potential. Within the *Enterobacteriaceae* DNA sequence space we found multiple associations of phages with multidrug resistant bacteria following whole genome sequencing of clinically-derived isolates. From a clinical perspective it would be valuable to understand if naturally-occuring phages impact disease outcome. The presence of lytic phage genomes in clinical isolates also hints at a reservoir of potential antimicrobial agents archived in strain collections. For the research community in general this is important as microbial cultures have previously been mined for such biological resources. Much time is spent within phage biology just accruing novel phages and often phages are searched for in environments where it is not known if they are present or not. This method of searching would require optimisation of activation but safe in the knowledge that the novel phages are present.

### Expanding Phage Diversity

Our study transforms the known diversity of phages and the discovery of over 100,000 complete lytic phage genomes within bacterial assemblies underscores the vast, largely unexplored phage diversity. This includes the significant expansion of the *Seoulvirus* family and the identification of new phage families such as the *Goslarvirus* and Bapsvirus clusters. These findings highlight the limitations of our current understanding of phage diversity and the potential for discovering new phages with unique genetic and biological properties. By systematically harvesting and cataloguing phages from diverse environmental and clinical samples, we can build a comprehensive library of phages. This expanded phage space not only enhances our understanding of phage biology and evolution but also provides a valuable resource for developing new phage-based applications in biotechnology and medicine.

### Future Directions

The unexpected discovery of lytic phages within bacterial assemblies opens several avenues for future research. Further studies are clearly needed to explore the mechanisms by which these phages associate with bacterial genomes without integrating into them. This will involve investigating the molecular interactions between phages and bacterial hosts, and identifying the environmental conditions that favour such associations. The fact that we have now this observation that phages used in therapy may have already been experienced by humans during infections, provides further evidence that phage therapy is not about subjecting humans to something ‘new’ but trying to understand the ecology enough to stack the odds in favour of the human rather than a bacterial pathogen. It will be highly relevant to see how such associations impact our understanding, development and deployment of phage therapy from both practical and regulatory standpoints.

The potential role of these phages in horizontal gene transfer, bacterial evolution and the ‘passing’ contribution to bacterial physiology should be explored. Also, by expanding the search to other bacterial species and environments will help determine the ubiquity and ecological reach of phages associated with bacteria in a quasi-stable way. As the control on such relationships is likely to be very finely tuned it will be important to determine how changing environmental parameters may alter the balance of bacterial-phage relationships, and how microbial ecosystems could shift in response to changing temperatures, pH and other altered physiological conditions.

Finally, of great practical use, leveraging new technologies within synthetic biology and cell-free systems will help translate these novel phages and to bring them to ‘life’, ultimately leading to new phages to be used in therapy or within agricultural, aquaculture or other applications to drive microbial ecosystems to a desired balance.

## Data Availability

XXX

## Methods

### BAPS

To identify complete lytic phage genomes within bacterial genome assemblies, we developed a comprehensive bioinformatic workflow, beginning with *assembly_summary_genbank*.*txt* containing assembly information from the NCBI database, downloaded on December 23, 2023. The initial step was to extract all bacterial and archaeal genomes represented by at least 50 distinct assemblies, resulting in 3,643,575 bacterial assemblies, spanning 1,226 bacteria/archaea species, totaling 230,974,966 contigs. We employed a stringent filtering approach, flagging contigs exceeding 1,000,000 base pairs (bp) as true bacteria/archaea(non-phage), and narrowed down to 114,681,711 contigs. Contigs exceeding 5,000 bp were considered potential phage candidates. Each remaining candidate contig was analysed using *phager*.*py*, a specialized machine learning tool designed to evaluate phage likelihood based on a set of biological features. Contigs scoring below 0.8 were excluded, refining our selection to those with a higher likelihood of phage origin, resulting in a set of 3,503,832 potential phage candidates.

Contigs were compared to NT reference database and annotated as bacterial if the sequence length of the mmseqs top hit exceeded 1,200,000 bp and the alignment length to the potential phage contig was larger than 2,100 bp, indicating potential bacterial origin. This resulted in 2,055,116 contigs annotated as “bacterial”. In addition to phage classification, plasmids were identified through database matches containing the word “plasmid,” resulting in 563,201annotations. All contigs were screened against a set of known phage databases, including INPHARED and selected PhageCloud sources (NCBI, HugePhages, Archeal Viruses, GPD, CHVD, GVDv1, and IMG/VR v4), where matches immediately suggested a lytic phage classification, resulting in 126,127 annotations. Furthermore, we identified phage clouds based on matches against the NCBI Nucleotide database^23^ containing the word ‘phage’ but not ‘prophage,’ excluding those overlapping with bacterial and temperate annotations. Clustering these phage candidates using Average Nucleotide Identity (ANI) resulted in 77,552 distinct clusters.

To differentiate between lytic and temperate phages, any cluster containing even a single integrase was labeled as temperate, resulting in 53,598 annotations. Contigs were classified as lytic if they exhibited a terminase large subunit hits (terL) while lacking integrase and transposase genes and had no anti-repressor proteins, resulting in 162,794 annotations. This workflow yielded 119,510 lytic phages, 602,285 plasmids, 146,575 temperate phages and 536,888 phage-like contigs where no certain decision could be made. The number of temperate phages or prophages is a great underestimate, as most of those had been removed in the very first screening of the 230 million contigs.

Table 1 from 1,226 organisms, to 3.6 million bacterial assemblies, to 230 million contigs to 3.5 million predicted phages to 119,510 lytic phages.

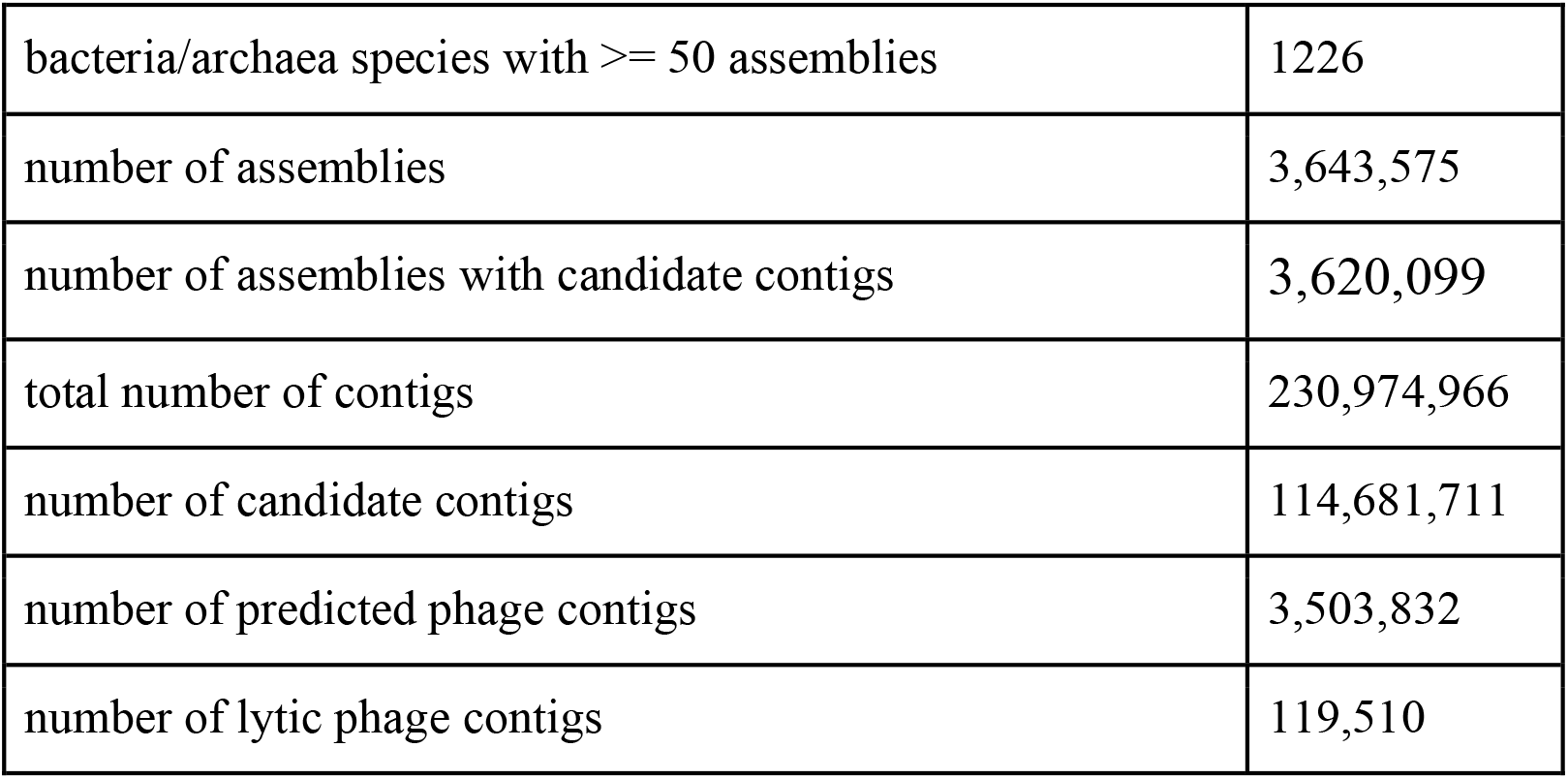

**Table 1.**

| Genome/Contig | Length (bp) | Accession Number | ANI with Munch (%) | Source | Location |
| --- | --- | --- | --- | --- | --- |
| Phage Munch | 350,103 | MK268344 | 100 | Cattle | Texas, USA |
| Phage PHA46 | 337,158 | PP191103 | 97.7 | River | Mexico |
| Phage SE-PL | 348,657 | MT001829 | 97.6 | Sewage | India |
| Phage 7t3 | 348,718 | MK773491 | 97.6 | Sewage | India |
| *S. enterica* PNUSAS388095 | 349,993 | ABNLLY010000004 | 99.5 | Unknown | PulseNet, USA |
| *S. enterica* PNUSAS432134 | 348,883 | ABRUYV010000006 | 98.3 | Unknown | PulseNet, USA |
| *S. enterica* PNUSAS219451 | 352,720 | ABBVBI010000007 | 98.4 | Intestine | PulseNet, USA |
| *S. enterica* OH-16-27174 | 349,812 | AAEKPK010000003 | 99 | Bovine faeces | GenomeTrakr, USA |
| *S. enterica* PNUSAS415804 | 343,049 | ABPXTM010000005 | 98.4 | Unknown | PulseNet, USA |
| *S. Infantis* 63028 | 350,071 | AAKURN010000002 | 99.3 | Human | PHE, UK |
| *S. Mbandaka* strain 110 | 350,137 | JAHSKT010000004 | 99.6 | Cattle | Texas, USA |

### Phager - development of a Phage-Likeliness Predictor

To efficiently screen millions of bacterial contigs for potential phage candidates, we developed Phager, a rapid phage-likeliness machine-learning predictor based on biological features. For training, we used a set of non-NCBI phage genomes from the PhageClouds database^24^ and bacterial genomes from the NCBI database. Initial data preparation involved gene prediction with Prodigal (v2.6.3)^25^ and the removal of potential prophage regions from the bacterial genomes. Prophages were identified using PhageBoost. Following this, rather than using complete bacterial genomes for training, we randomly selected bacterial genome fragments whose lengths fall within the range of typical phage genome sizes. This approach was chosen to reduce the marked size differences between bacterial and phage genomes.

Next, we calculated biological features for each gene in these genomes, following the methodologies established in PhageBoost and PhageLeads^26,27^. Each genome was then segmented into overlapping gene triplets along a shifting reading frame, with their associated features. Instead of using entire genome sequences, each gene feature triplet served as an individual training unit. The model was trained using LightGBM^28^.

To ensure accuracy and reliability, we rigorously evaluated the predictor using all known phage genomes in the NCBI database. Phager showed high precision in distinguishing phage contigs from bacterial ones, making it a fast and robust tool for large-scale genomic screenings. This approach allowed us to systematically identify and categorise potential lytic phage candidates from a vast dataset, greatly improving our ability to explore phage diversity within bacterial genome assemblies.

### Pairwise genome distance estimation using BinDash (Between phages and BAPS)

A total of 66 publicly available lytic phage genomes previously used in phage therapy trials were selected and screened against the BAPS dataset to identify candidate contigs with similar genetic content (see Table 3). Genomic similarity between phage genomes and BAPS contigs was estimated using BinDash^29^ (version 2.1). Two genome distance thresholds were applied to classify levels of similarity: a threshold of ≤ 0.2 was used to identify closely associated contigs, while a more stringent threshold of ≤ 0.05 was used to define strongly associated contigs with high sequence similarity. Matches were recorded only when one or more BAPS contigs fell within the respective distance thresholds.

**Table 2.** Analysis of read mapping statistics for *Goslarvirus* BAPS and host bacteria.

| Goslarviruses | Genome Size (bp) | Total reads | Reads Mapped to BAPS contig (%) | Unmapped reads to host (%) |
| --- | --- | --- | --- | --- |
| *S_Heidelberg_57606* | 233,528 | 2,965,546 | 55,142 (1.86%) | 2,910,404 (98.14%) |
| *S_enterica_PNUSAS134354* | 234,406 | 2,108,438 | 96,329 (4.57%) | 2,012,109 (95.43%) |
| *S_Corvallis_69921* | 235,022 | 4,591,164 | 1,384,765 (30.16%) | 3,206,399 (69.84%) |
| *Ecoli_PNUSAE136084* | 235,174 | 2,307,558 | 133,274 (5.78%) | 2,174,284 (94.22%) |
| *S_enterica_PNUSAS313451* | 235,335 | 2,028,144 | 286,553 (14.13%) | 1,741,591 (85.87%) |
| *Ecoli_92_1R* | 235,435 | 7,242,968 | 3,324,926 (45.91%) | 3,918,042 (54.09%) |
| *S_flexneri_PNUSAE097064* | 235,676 | 1,453,770 | 130,182 (8.95%) | 1,323,588 (91.05%) |
| *E_hormaechei_SAMEA112238924* | 235,745 | 1,050,316 | 41,328 (3.93%) | 1,008,988 (96.07%) |
| *S_enterica_PNUSAS242309* | 235,997 | 1,496,512 | 40,271 (2.69%) | 1,456,241 (97.31%) |
| *Ecoli_G64_1R* | 236,079 | 6,661,988 | 126,637 (1.90%) | 6,535,351 (98.10%) |
| *S_Enteritidis_123963* | 236,357 | 2,755,372 | 993,941 (36.07%) | 1,761,431 (63.93%) |
| *Ecoli_PSU_2039* | 236,559 | 1,471,050 | 44,720 (3.04%) | 1,426,330 (96.96%) |
| *S_enterica_45561* | 236,560 | 2,406,508 | 122,132 (5.08%) | 2,284,376 (94.92%) |
| *S_enterica*_PNUSAS226980 | 236,597 | 2,459,670 | 46,988 (1.91%) | 2,412,682 (98.09%) |
| *S_flexneri*_PNUSAE015436 | 236,870 | 2,215,162 | 2,137,243 (96.48%) | 77,919 (3.52%) |
| *S_Newport*_1675657 | 237,051 | 8,372,828 | 493,179 (5.89%) | 7,879,649 (94.11%) |
| *S_enterica*_PNUSAS179791 | 237,225 | 2,166,716 | 93,358 (4.31%) | 2,073,358 (95.69%) |
| Goslar | 237,307 | N/A | N/A | N/A |
| *S_Newport*_1343546 | 237,553 | 2,975,760 | 2,951,938 (99.20%) | 23,822 (0.80%) |
| *S_Enteritidis*_227002 | 237,669 | 3,252,092 | 365,990 (11.25%) | 2,886,102 (88.75%) |
| *Ecoli*_ML5_10A | 238,122 | 6,167,110 | 804,421 (13.04%) | 5,362,689 (86.96%) |
| *S_Enteritidis*_224060 | 238,368 | 3,557,196 | 113,295 (3.18%) | 3,443,901 (96.82%) |
| *Ecoli*_O157_RM10569 | 238,408 | 1,658,616 | 26,815 (1.62%) | 1,631,801 (98.38%) |
| *Ecoli*_O157_RM11324 | 238,410 | 1,420,568 | 42,381 (2.98%) | 1,378,187 (97.02%) |
| *Ecoli*_O157_RM11325 | 238,410 | 1,661,394 | 108,239 (6.51%) | 1,553,155 (93.49%) |
| *Ecoli*_P59_1R | 238,477 | 13,154,236 | 722,271 (5.49%) | 12,431,965 (94.51%) |
| *Ecoli*_1729750 | 238,662 | 6,536,282 | 161,912 (2.48%) | 6,374,370 (97.52%) |
| *S_enterica*_PNUSAS285227 | 238,746 | 1,201,978 | 1,183,976 (98.50%) | 18,002 (1.50%) |
| *S_flexneri*_PNUSAE118324 | 238,774 | 3,523,916 | 1,737,237 (49.3%) | 1,786,679 (50.7%) |
| *Ecoli*_SAMEA11308066 | 238,969 | 8,715,550 | 3,499,478 (40.15%) | 5,216,072 (59.85%) |
| *Ecoli*_PNUSAE008133 | 239,119 | 2,313,434 | 825,008 (35.66%) | 1,488,426 (64.34%) |
| *S_enterica*_PNUSAS294194 | 239,153 | 3,981,692 | 24,387 (0.61%) | 3,957,305 (99.39%) |
| *S_Kingston*_285832 | 239,299 | 2,910,890 | 164,906 (5.67%) | 2,745,984 (94.33%) |
| *Ecoli*_CFSAN086594 | 239,345 | 851,272 | 177,935 (20.90%) | 673,337 (79.10%) |
| *S_enterica*_37067 | 239,430 | 1,802,014 | 128,790 (7.15%) | 1,673,224 (92.85%) |
| *S_Bovismorbificans*_ADRDL_16_12667 | 239,661 | 1,450,502 | 18,378 (1.27%) | 1,432,124 (98.73%) |
| *Ecoli*_O157_RM10577 | 239,682 | 1,994,212 | 62,635 (3.14%) | 1,931,577 (96.86%) |
| *S_Enteritidis*_216974 | 239,886 | 3,781,446 | 1,436,553 (37.99%) | 2,344,893 (62.01%) |
| *S_Kentucky*_36997 | 240,786 | 4,345,736 | 138,410 (3.18%) | 4,207,326 (96.82%) |
| *Ecoli*_P17_2R | 240,923 | 11,469,310 | 1,107,960 (9.66%) | 10,361,350 (90.34%) |
| *S_Corvallis*_37805 | 241,301 | 2,145,282 | 173,395 (8.08%) | 1,971,887 (91.92%) |
| *S_Enteritidis*_597045 | 242,403 | 5,455,338 | 33,552 (0.62%) | 5,421,786 (99.38%) |
| *S_Enteritidis*_118870 | 242,647 | 3,556,032 | 1,139,505 (32.04%) | 2,416,527 (67.96%) |
| *S_Typhimurium*_404312 | 243,202 | 2,205,724 | 77,644 (3.52%) | 2,128,080 (96.48%) |
| *S_sonnei*_PNUSAE090274 | 243,941 | 1,426,540 | 33,689 (2.36%) | 1,392,851 (97.64%) |
| *S_Newport*_365400 | 243,993 | 3,478,588 | 144,023 (4.14%) | 3,334,565 (95.86%) |
| *S_Panama*_161075 | 247,237 | 6,340,498 | 2,898,622 (45.72%) | 3,441,876 (54.28%) |
| *S_flexneri*_PNUSAE151852 | 247,544 | 1,349,274 | 119,759 (8.88%) | 1,229,515 (91.12%) |
| *Ecoli*_O157_RM5667 | 248,066 | 4,995,936 | 441,142 (8.83%) | 4,554,794 (91.17%) |
| *Ecoli*_O157_RM7446 | 248,130 | 3,970,208 | 179,883 (4.53%) | 3,790,325 (95.47%) |
| *Ecoli*_O157_RM5672 | 248,132 | 3,266,840 | 343,184 (10.51%) | 2,923,656 (89.49%) |
| *Ecoli*_O157_RM13637 | 248,886 | 1,268,536 | 49,339 (3.89%) | 1,219,197 (96.11%) |
| *S_Enteritidis*_FD01851371_1 | 236,335 | N/A | N/A | N/A |
| *Ecoli*_Q0226 | 240,999 | N/A | N/A | N/A |
| *Ecoli_IN_MR364R1* | 237,721 | N/A | N/A | N/A |
| *Ecoli_IN_MR364S3* | 237,848 | N/A | N/A | N/A |

**Table 3.** Analysis of phages associated with phage therapeutic studies reveals an abundance of BAPS contigs in pathogen genome assemblies.

| Accession No. | Phage | Genome size (bp) | Host strain | Found in BAPS | BAPS under distance threshold $\leq 0.2$ | BAPS under distance threshold $\leq 0.05$ |
| --- | --- | --- | --- | --- | --- | --- |
| MT949699 | Maestro | 169176 | *A. baumannii*<sup>39</sup> | Yes | 1 | 1 |
| OL770258 | AB-Navy1 | 166113 | *A. baumannii*<sup>39</sup> | Yes | 1 | 0 |
| OL770259 | AB-Navy4 | 166964 | *A. baumannii*<sup>39</sup> | Yes | 1 | 0 |
| OL770260 | AB-Navy71 | 166382 | *A. baumannii*<sup>39</sup> | Yes | 1 | 1 |
| OL770261 | AB-Navy97 | 165480 | *A. baumannii*<sup>39</sup> | Yes | 1 | 0 |
| OL770262 | AC4 | 168186 | *A. baumannii*<sup>39</sup> | Yes | 1 | 0 |
| OL770263 | AbTP3phi1 | 42093 | *A. baumannii*<sup>39</sup> | Yes | 38 | 0 |
| MK278859 | AbKT21phiIII | 40898 | *A. baumannii*<sup>16</sup> | Yes | 38 | 0 |
| NC_025462 | Acibel004 | 99730 | *A. baumannii*<sup>17</sup> | No | - | - |
| NC_025457 | Acibel007 | 42654 | *A. baumannii*<sup>17</sup> | No | - | - |
| NC_023556 | JWAlpha | 72329 | *A. xylosoxidans*<sup>17</sup> | No | - | - |
| KF787094 | JWDelta | 73659 | *A. xylosoxidans*<sup>17</sup> | No | - | - |
| OQ938574 | JWT | 46410 | *A. xylosoxidans*<sup>17</sup> | Yes | 1 | 1 |
| OQ974181 | 2-1 | 82711 | *A. xylosoxidans*<sup>17</sup> | No | - | - |
| NC_028768 | JWX | 49714 | *A. xylosoxidans*<sup>17</sup> | Yes | 1 | 1 |
| OQ116603 | UZM3 | 46054 | *B. fragilis*<sup>17</sup> | Yes | 4 | 0 |
| MW883060 | vB_EcoP_SYGE1 | 39713 | *E. coli*<sup>40</sup> | Yes | 5 | 0 |
| MW883061 | vB_EcoM_SYGMH1 | 137423 | *E. coli*<sup>40</sup> | Yes | 118 | 4 |
| MW883059 | vB_EcoM_SYGD1 | 171255 | *E. coli*<sup>40</sup> | Yes | 166 | 60 |
| OL870317 | E4 | 169163 | *E. coli*<sup>40</sup> | Yes | 159 | 62 |
| PP355091 | T4 | 167904 | *E. coli*<sup>41</sup> | Yes | 170 | 60 |
| MZ333457 | SSsP-1 | 57270 | *E. faecium*<sup>42</sup> | Yes | 1 | 0 |
| MZ333462 | GVEsP-1 | 149913 | *E. faecium*<sup>42</sup> | Yes | 1 | 0 |
| MW004544 | EFGrKN | 147532 | *Enterococcus spp.*<sup>17</sup> | Yes | 1 | 0 |
| MW004545 | EFGrNG | 145199 | *Enterococcus spp.*<sup>17</sup> | Yes | 1 | 0 |
| OL870612 | Efs7 | 56144 | *Enterococcus spp.*<sup>17</sup> | Yes | 1 | 0 |
| NC_048143 | KpKT21phi1 | 49106 | *K. pneumoniae*<sup>16</sup> | Yes | 17 | 8 |
| MW448170 | M1 | 176329 | *K. pneumoniae*<sup>17</sup> | Yes | 7 | 4 |
| DQ004855 | P100 | 131384 | *L. monocytogenes*<sup>43</sup> | No | - | - |
| OQ988004 | 8UZL | 47806 | *M. abscessus*<sup>17</sup> | No | - | - |
| OP875100 | SPA01 | 93536 | *P. aeruginosa*<sup>44</sup> | Yes | 2 | 0 |
| OP875101 | SPA05 | 93656 | *P. aeruginosa*<sup>44</sup> | Yes | 2 | 2 |
| OM870967 | Ps12on-D | 88705 | *P. aeruginosa*<sup>45</sup> | Yes | 2 | 2 |
| OM870968 | Ps25 | 87887 | *P. aeruginosa*<sup>45</sup> | Yes | 2 | 2 |
| OM870969 | PsCh | 92710 | *P. aeruginosa*<sup>45</sup> | Yes | 2 | 2 |
| OM870970 | PsIn | 92515 | *P. aeruginosa*<sup>45</sup> | Yes | 2 | 2 |
| ON631220 | OMKO1 | 281755 | *P. aeruginosa*<sup>46</sup> | Yes | 11 | 2 |
| HM624080 | PA1Ø | 34553 | *P. aeruginosa*<sup>47</sup> | Yes | 3146 | 462 |
| NC_011703 | 14-1 | 66235 | *P. aeruginosa*<sup>17</sup> | Yes | 8 | 4 |
| OP292288 | PNM | 42721 | *P. aeruginosa*<sup>17</sup> | No | - | - |
| OQ850183 | PT07 | 94660 | *P. aeruginosa*<sup>17</sup> | Yes | 2 | 2 |
| ON815901 | 4029 | 72063 | *P. aeruginosa*<sup>17</sup> | Yes | 1 | 1 |
| ON815902 | 4032 | 72063 | *P. aeruginosa*<sup>17</sup> | Yes | 1 | 1 |
| ON815903 | 4034 | 72063 | *P. aeruginosa*<sup>17</sup> | Yes | 1 | 1 |
| OQ872152 | 4P | 66197 | *P. aeruginosa*<sup>17</sup> | Yes | 8 | 4 |
| NC_041870 | DP1 | 66158 | *P. aeruginosa*<sup>17</sup> | Yes | 8 | 5 |
| OQ925957 | Phage C | 65622 | *P. aeruginosa*<sup>17</sup> | Yes | 8 | 3 |
| OL741432 | Isf-Pm2 | 167727 | *P. mirabilis*<sup>12</sup> | Yes | 164 | 62 |
| OL741431 | Isf-Pm1 | 58354 | *P. mirabilis*<sup>12</sup> | Yes | 5 | 5 |
| MT080595 | vB_SauM_EW41 | 132999 | *S. aureus*<sup>48</sup> | Yes | 0 | 0 |
| OQ129426 | vB_SauM_JDF86 | 146296 | *S. aureus*<sup>40</sup> | Yes | 0 | 0 |
| MK417516 | J-Sa36 | 148312 | *S. aureus*<sup>49,50</sup> | Yes | 14 | 3 |
| MK417515 | Sa83 | 149108 | *S. aureus*<sup>49,50</sup> | Yes | 14 | 3 |
| MK417514 | Sa87 | 146004 | *S. aureus*<sup>49,50</sup> | Yes | 14 | 3 |
| KF766114 | Phage_K | 148317 | *S. aureus*<sup>13,14</sup> | Yes | 14 | 3 |
| OQ129425 | vB_SauM_JDYN | 143839 | *S. aureus*<sup>40</sup> | Yes | 6 | 0 |
| NC_023009 | Sb-1 | 127188 | *S. aureus*<sup>15,51,52</sup> | Yes | 12 | 3 |
| NC_047720 | ISP | 138339 | *S. aureus*<sup>17</sup> | Yes | 14 | 3 |
| MT596503 | BE06 | 140659 | *S. aureus*<sup>17</sup> | Yes | 2 | 1 |
| OM735686 | BUCT700 | 43214 | *S. maltophilia*<sup>17</sup> | No | - | - |
| KX130864 | SHBML-50-1 | 166634 | *S. sonnei*<sup>11</sup> | Yes | 166 | 61 |
| KX130863 | SHSML-45 | 108050 | *S. sonnei*<sup>11</sup> | Yes | 44 | 0 |
| KX130862 | SHFML-26 | 168993 | *S. sonnei*<sup>11</sup> | Yes | 164 | 61 |
| KX130861 | SHFML-11 | 170650 | *S. sonnei*<sup>11</sup> | Yes | 160 | 60 |
| KX130865 | SHSML-52-1 | 169621 | *S. sonnei*<sup>11</sup> | Yes | 77 | 15 |
| OQ680630 | PG288 | 76120 | *V. parahaemolyticus*<sup>53</sup> | Yes | 7 | 4 |

### Environmental BAPS in human microbiomes

To examine the prevalence and distribution of lytic BAPS phage genomes in human microbiomes, we analysed five large metagenomic datasets. Four of these were recently published, phage-filtered metagenomic collections. The Early Life Gut Virome (ELGV) dataset comprises a catalog of 160,478 non-redundant viral sequences identified during the first three years of life ^30^. The Gut Virome Database (GVDv1) contains 33,242 unique viral populations classified at the species level ^31^. The Gut Phage Database (GPD) includes approximately 142,000 non-redundant viral genomes recovered from a global dataset of 28,060 gut metagenomes and 2,898 reference bacterial genomes ^32^. The Metagenomic Gut Virus (MGV) catalog consists of 189,680 viral genomes derived from 11,810 stool metagenomes ^33^. In addition to the four virome collections, we included the unfiltered whole-genome metagenomic sequencing data from the NIH Human Microbiome Project (HMP). The dataset comprised 3,778 samples collected from healthy individuals across diverse anatomical sites, including the oral cavity, nasal region, skin, gastrointestinal tract (feces), throat, and female reproductive tract ^34^. All five metagenomic collections were compared against the 130,805 predicted lytic phages that were identified in the BAPS using BLASTN v2.16.0 ^35^. Hits were filtered using a threshold of 90% identity and 500 bp minimum alignment length. Total non-overlapping match length was calculated in order to summarise matches between the identical sequences with different genomic locations. Focusing on *Felixounavirus* and jumbo phages, the final datasets contained the BAPS-metagenome matches of at least 80,000 bp. ANI was also calculated to showcase the level of similarity among the sequences.

### Taxonomic validation of BAPS assemblies via reference BLAST and domain consistency analysis

To validate the taxonomic assignments of BAPS-derived assemblies, we extracted the longest scaffold from each matched reference genome and performed BLASTN searches (NCBI nt database, Feb 2024 release) against this sequence using up to 5 top hits per scaffold (- max_target_seqs 5, -max_hsps 1, -evalue 1e-4). BLAST results were parsed to retrieve taxonomic identifiers, which were then compared to the BAPS-assigned taxonomy using the ETE3 toolkit and a local NCBI taxonomy SQLite database. We determined the lowest common rank between the assigned and BLAST-derived taxids, evaluated domain-level consistency (e.g., Bacteria vs. Viruses), and provided an interpretative reasoning string for each comparison. For every BAPS contig, we included all five BLAST hits from the corresponding reference scaffold in a merged summary table, while also producing filtered outputs containing only the top hit per contig and a subset of contigs with mismatched taxonomic domains. Contigs with no significant BLAST hit were retained and flagged, ensuring a complete overview of classification confidence across all ∼130,000 BAPS assemblies.

### Taxonomic classification of BAPS

To classify selected phage sequences at the genus and species level, *taxMyPhage* was used to first classify all contigs into existing genera and species using the “run” option ^8^. Contigs not classified into current ICTV genera or species from PhageClouds clusters, were then analysed with the “similarity” option to identify new genera and species based on ICTV standards ^36^. In order to not overestimate the number of new phage species, only contigs that had a length that was 90% similarity to the closest isolated phage genome were used to determine the number of species. Further classification of the phage genomes at higher taxonomic levels was achieved using the standalone version of ViPTreeGen, with default settings ^37^, trees were viewed in iTOL v7 ^38^

## Notes

### Competing Interest Statement

The authors have declared no competing interest.

